# Computational design of anti-CRISPR proteins with improved inhibition potency and expanded specificity

**DOI:** 10.1101/685032

**Authors:** Jan Mathony, Zander Harteveld, Carolin Schmelas, Julius Upmeier zu Belzen, Sabine Aschenbrenner, Mareike D. Hoffmann, Christina Stengl, Andreas Scheck, Stéphane Rosset, Dirk Grimm, Roland Eils, Bruno E. Correia, Dominik Niopek

**Author notes:** These authors contributed equally to this work (Jan Mathony, Zander Harteveld, Carolin Schmelas). To whom correspondence should be addressed Bruno E. Correia, EPFL IBI STI LPDI Station 19, CH-1015 Lausanne, Switzerland, Dominik Niopek, Im Neuenheimer Feld 267, 69120 Heidelberg, Germany.

## Abstract

Anti-CRISPR (Acr) proteins are bacteriophage-derived antagonists of CRISPR-Cas systems. To date, Acrs were obtained either by mining sequence databanks or experimentally screening phage collections, both of which yield a limited repertoire of naturally occurring variants. Here, we applied structure-based engineering on AcrIIC1, a broad-spectrum inhibitor of type II-C CRISPR systems, to improve its efficacy and expand its specificity. We first show that fusing exogenous protein domains into AcrIIC1 dramatically enhances inhibition of the natural *Neisseria meningitidis* Cas9 target. Then, using structure-guided design, we converted AcrIIC1 into AcrX, a potent inhibitor of the type II-A CRISPR-Cas9 from *Staphylococcus aureus* widely applied for *in vivo* genome editing. Our work introduces designer Acrs as important biotechnological tools and provides an innovative strategy to safeguard the CRISPR technology.

---

The detailed characterization of bacterial CRISPR-Cas systems^1^ and their adaptation for precise genome engineering in mammalian cells^2, 3^ has revolutionized the life sciences and enabled novel applications in biotechnology and medicine. The recent discovery of phage-derived anti-CRISPR proteins^4-6^, *i.e.* potent inhibitors of Cas effectors, provides a shut-off mechanism that can keep this powerful technology in check^7^ and enhance the precision at which genome perturbations can be made^8-11^. While mining of sequence databases and screening of phage libraries proved to be powerful strategies to discover Acrs targeting a variety of Cas effectors^5, 6, 12-18^, these approaches are inherently limited to the naturally occurring protein sequence space. Moreover, for various Cas effectors of major biotechnological interest, nature might be lacking (efficient) anti-CRISPR counterparts.

In this work, we sought to apply protein engineering to design artificial Acrs that exhibit enhanced inhibition potency and expanded target specificity (Fig. 1). As starting point, we used AcrIIC1^14^, a broad-spectrum inhibitor targeting various type II-C Cas9s, including those from *Neisseria meningitidis* (*Nme*), *Geobacillus stearothermophilus* and *Campylobacter jejuni.* AcrIIC1 binds the conserved catalytic HNH domain and locks Cas9 in a DNA binding-competent, but catalytically inactive state. This unique inhibitory mechanism might explain why AcrIIC1 is a rather weak inhibitor^15^ as compared to its related proteins AcrIIC3, -C4 and -C5, all of which interfere with Cas9 DNA binding. Importantly, biochemical assays suggest tight binding of AcrIIC1 to the *Nme*Cas9 HNH domain (K_D_ ∼ 6.3 nM (Ref. 2)), indicating that it is the inhibitory mechanism rather than the affinity which leads to the suboptimal performance of AcrIIC1.

**Fig. 1.**
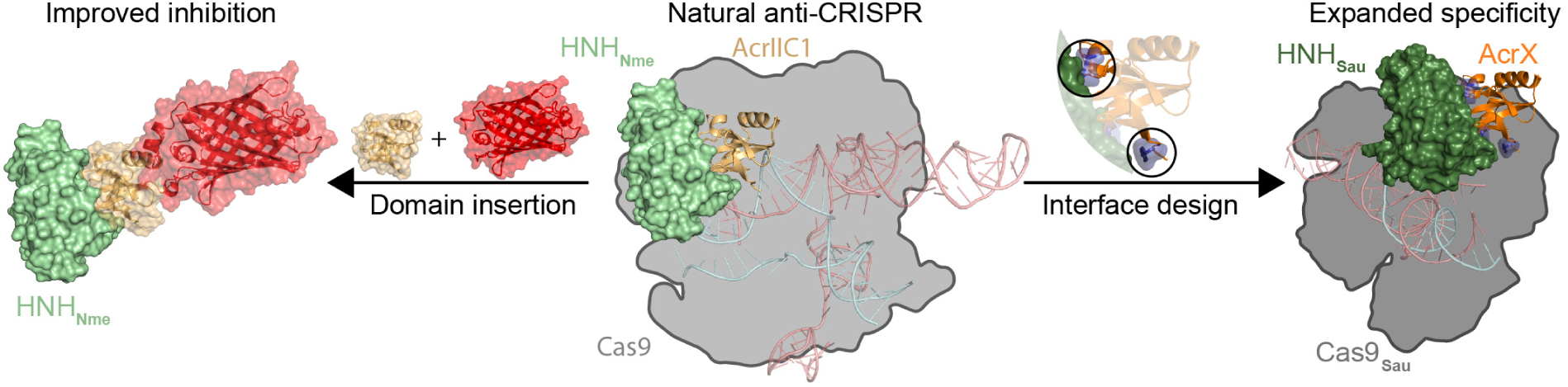
Improving potency and expanding specificity of anti-CRISPR proteins. Domain insertion into AcrIIC1 loop 5 yields chimeric inhibitors with improved inhibition potency, while structure-based engineering of its HNH-binding interface expands its target specificity (PDB 4ZIN, 5VGB, 5F9R and 5CZZ).

We hypothesized that inserting a sufficiently large, exogenous domain into AcrIIC1 could result in an inhibitor that would still bind tightly to the HNH domain, but – on top of inhibiting Cas9 catalytic activity – could block additional, important mechanistic steps such as DNA binding (Fig. 1, left). To test this hypothesis, we first created a structural model of the AcrIIC1-Cas9-sgRNA complex (see Methods for details) and interrogated it for AcrIIC1 surface sites that could be amenable for engineering. AcrIIC1 loop 5 appeared to be an ideal candidate (Supplementary Fig. 1), as it is located close to the DNA binding groove when assuming the HNH-docked Cas9 state^19^, but distal from the HNH-interacting surface necessary for activity that we wished to preserve. We then created 11 different AcrIIC1 domain fusions which carry either an mCherry (∼27 kDa), *Avena sativa* LOV2 (∼17 kDa) or PDZ domain (∼9 kDa) at different positions in loop 5 and optional, flanking GS-linkers or short deletions (Supplementary Fig. 2a). These chimeric Acrs were screened by T7 endonuclease assay for their ability to inhibit *Nme*Cas9 cleavage of the IL2RG locus in HEK 293T cells (Supplementary Fig. 2b). Remarkably, several AcrIIC1-LOV2 and AcrIIC1-mCherry chimeras mediated highly potent *Nme*Cas9 inhibition largely exceeding that of the parent AcrIIC1 (Fig. 2a,b, Supplementary Figs. 2b and 3). The AcrIIC1-mCherry chimera #10 (GSG-mCherry-GSG inserted between AcrIIC1 residues Y70 and A71; Supplementary Fig. 2a) showed an improvement superior to 12-fold in *Nme*Cas9 inhibition as compared to wild-type AcrIIC1 on the four tested loci (Fig. 2a and b), and performed equivalently or better than AcrIIC3, so-far the most potent *Nme*Cas9 inhibitor in mammalian cells^14^ (Fig. 2b).

**Fig. 2.**
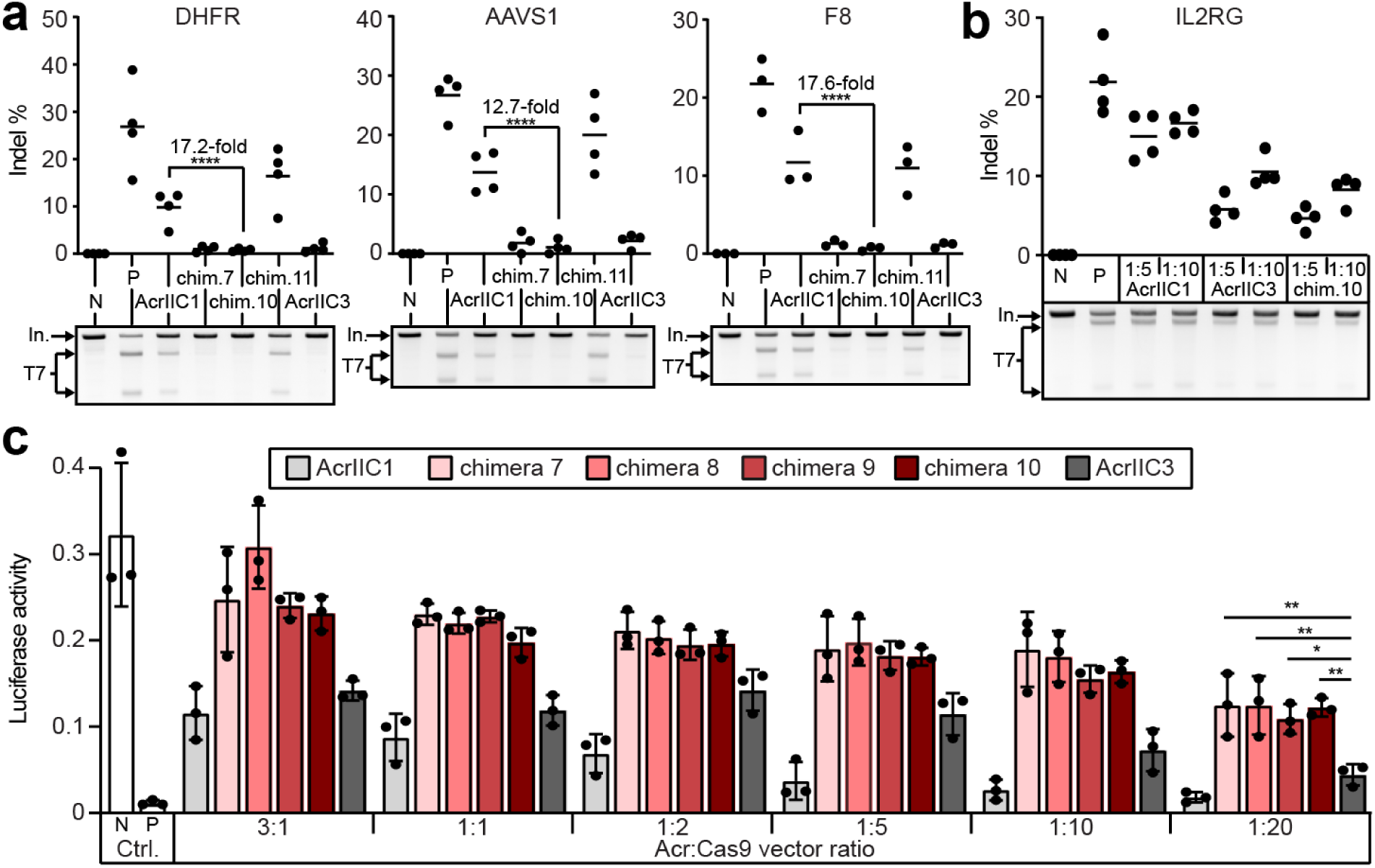
Domain insertion into AcrIIC1 yields a highly potent *Nme*Cas9 inhibitor. (**a**,**b**) HEK 293T cells were co-transfected with vectors expressing *Nme*Cas9, the indicated Acr and sgRNAs targeting different genomic loci followed by T7 endonuclease assay. In **a**, Acr:*Nme*Cas9 vector ratio used during transfection was 1:1, while in **b**, the indicated, low Acr:*Nme*Cas9 vector ratios were used. Representative T7 gel images and corresponding quantification of indel frequencies are shown. Lines in plots show means, dots are individual data points for n = 4 (DHFR, AAVS1 and IL2RG locus) or n = 3 (F8 locus) independent experiments. Chim., AcrIIC1-mCherry chimeras in Supplementary Fig. 2. In., input band. T7, T7 cleavage fragments. (**c**) AcrIIC1-mCherry chimeras outperform wild-type AcrIIC1 and AcrIIC3. Cells were co-transfected with vectors encoding *Nme*Cas9, a firefly luciferase reporter and corresponding reporter-targeting sgRNA as well as the indicated Acr followed by luciferase assay. Bars indicate means, error bars the SD for n = 3 independent experiments. (**a-c**) N, negative (Cas 9 only (**a**,**b**) or reporter only (**c**) control (Ctrl.)). P, positive control (Cas9 + sgRNA (**a**,**b**) or reporter + Cas9 (**c**)). \**P < 0.05*, \*\**P < 0.01*, \*\*\**P < 0.001*, \*\*\*\**P < 0.0001* by one-way ANOVA with Bonferroni correction.

To further characterize the gain in inhibition, we employed a reporter assay in which *Nme*Cas9 cleaves a firefly luciferase transgene, thereby resulting in luciferase knockout. We co-transfected cells with the reporter, *Nme*Cas9 and either (i) the parent AcrIIC1, (ii) the AcrIIC3 benchmark or (iii) different, engineered AcrIIC1-mCherry chimeras. During transfection, we varied the Acr:Cas9 vector ratios from 3:1 to 1:20. The chimeric inhibitors outperformed both, wild-type AcrIIC1 as well as AcrIIC3 and showed potent Cas9 inhibition even at very low Acr:Cas9 vector ratios (Fig. 2c).

To elucidate this particularly efficient inhibitory mechanism, we performed an assay in which the firefly luciferase gene was targeted by a *Nme*Cas9 nickase carrying an impaired RuvC, but a wild-type HNH domain (see Supplementary Note for details). Luciferase activity was potently rescued in the presence of the AcrIIC1-mCherry chimeric inhibitor, comparable to the AcrIIC3 benchmark (Supplementary Fig. 4). Notably, AcrIIC1 also mediated some rescue of luciferase activity, reaching precisely the level observed when targeting the same reporter with a catalytically dead (*d*)*Nme*Cas9 which inhibits firefly luciferase expression exclusively via CRISPR interference (Supplementary Fig. 4). This suggests that on top of blocking the catalytic activity (akin to native AcrIIC1), the chimeric inhibitors also interfere - at least to some extent - with *Nme*Cas9 DNA binding (see Supplementary Note). Together, these experiments demonstrate that domain insertion can yield anti-CRISPR proteins with far superior inhibition potency than that of natural anti-CRISPRs, thereby enabling extremely tight control of Cas9.

Having shown that AcrIIC1 could be modified to enhance its potency, we next aimed to expand its specificity. The Cas9 from *Staphylococcus aureus* is a type II-A CRISPR effector widely employed for *in vivo* genome editing^20^. Due to its favorable, small size (3.2 kb), *Staphylococcus aureus* (*Sau*)Cas9 can easily be packaged into Adeno-associated virus (AAV) particles^20^, which are prime vector candidates for therapeutic CRISPR applications^21, 22^. Importantly, no anti-CRISPR proteins have yet been discovered that would allow blocking *Sau*Cas9 activity. We speculated that AcrIIC1 might represent an ideal starting point to engineer an artificial *Sau*Cas9 inhibitor, as the overall fold of the *Sau*Cas9 HNH domain is similar to that of *Nme*Cas9^19^, albeit substantial differences exist at the sequence level (sequence identity is only 33.7 %, Supplementary Fig. 5). Recent *in vitro* data suggest that wild-type AcrIIC1 fails to inhibit *Sau*Cas9 function^23^. To independently confirm this finding, we co-expressed AcrIIC1 in HEK 293T cells together with *Sau*Cas9 and sgRNAs targeting different loci and performed T7 endonuclease assays to detect formation of insertions and deletions (indels). Strikingly, we observed a reproducible, albeit rather mild reduction of indels in the AcrIIC1 samples to about 50 % of the positive controls (Supplementary Fig. 6), indicating that AcrIIC1 can also bind the *Sau*Cas9 HNH domain, though likely with a compromised affinity.

Based on this assumption we sought to enhance *Sau*Cas9 binding. We generated a structural model of *Sau*Cas9 HNH domain in complex with AcrIIC1 and investigated the differences in the AcrIIC1 interacting surface as compared to the *Nme*Cas9 HNH (Fig. 3). Two regions in *Sau*Cas9 HNH domain showed suboptimal contacts to corresponding AcrIIC1 residues (Fig. 3). We then performed per-residue *in silico* mutagenesis using Rosetta design^24^ followed by manual inspection, which suggested ten AcrIIC1 candidate mutations to improve binding to the *Sau*Cas9 HNH domain (Supplementary Figs. 7 and 8). We tested these mutants, first individually, in genome editing experiments targeting the EMX1 locus and then iteratively combined the most promising variants in subsequent screening rounds (Supplementary Fig. 9). Of note, we decreased the Acr:Cas9 ratio with each screening round to better resolve the performance of improving candidates. After only three rounds, we arrived at a triple mutant (N3F, D15Q, A48I; Fig. 3 and Supplementary Fig. 10), which we confirmed to be well folded in solution (Supplementary Fig. 11) and which achieved a near-complete blocking of EMX1 editing by *Sau*Cas9 (Supplementary Fig. 9). Remarkably, this mutant also retained *Nme*Cas9 inhibitory activity comparable to wild-type AcrIIC1 (Supplementary Fig. 12, *Nme*Cas9 samples), indicating that our engineering approach had not just re-targeted AcrIIC1 to *Sau*Cas9, but rather expanded its specificity towards this orthologue. We therefore named the mutant AcrX for its ability to target both, type II-C and II-A CRISPR effectors, which – to our best knowledge – has not yet been observed for any natural Acr.

**Fig. 3.**
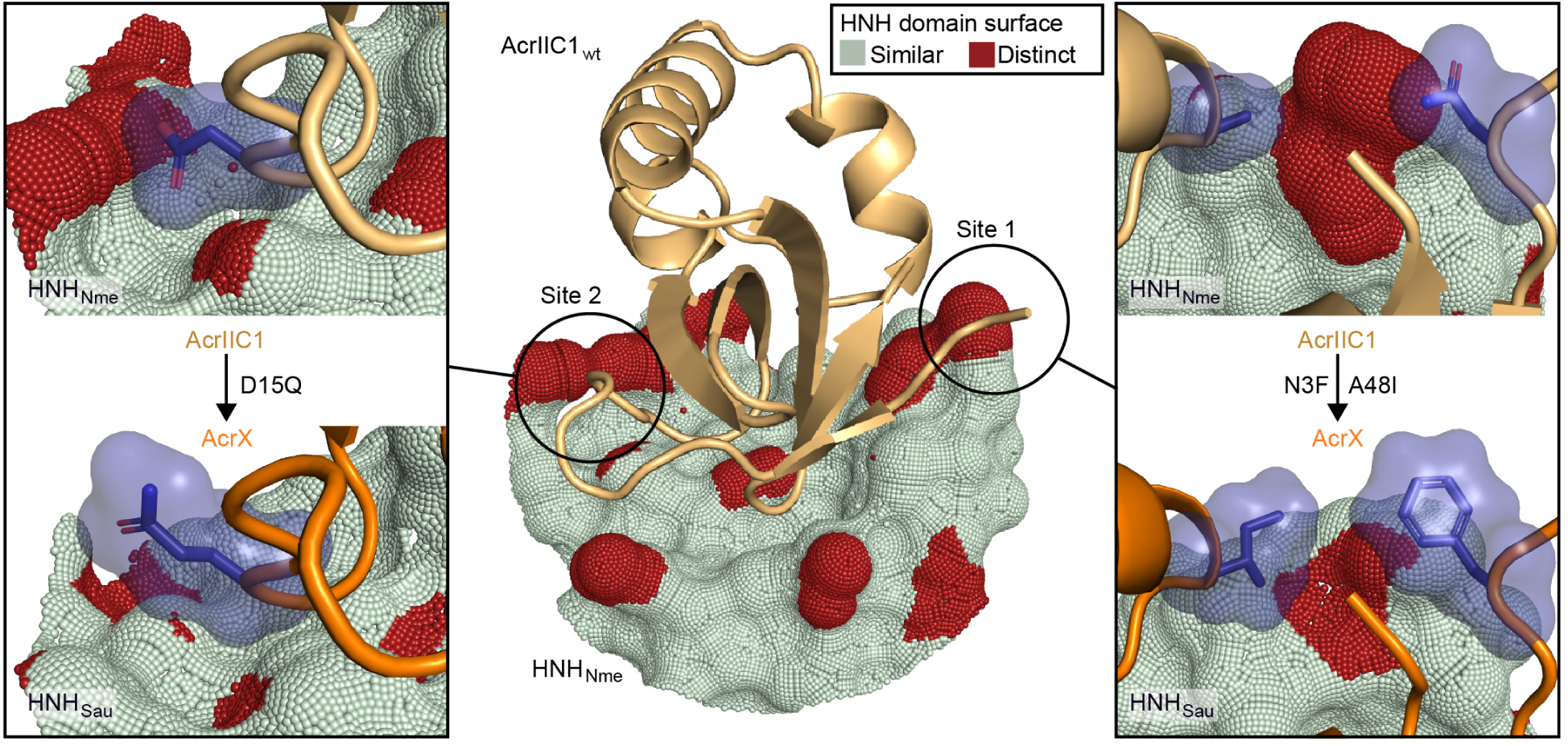
Structure-guided design of AcrX, an anti-CRISPR protein targeting *S. aureus* Cas9. Center: Structure showing AcrIIC1 binding to the *Nme*Cas9 HNH domain surface. Red patches indicate regions at which the *Sau*Cas9 HNH surface displays deviations of at least 1 Å as compared to the *Nme*Cas9 HNH surface, highlighting the most important sites to target by mutagenesis. Left/right: Comparison of wild-type AcrIIC1 residues (D15 (left), N3 and A48 (right)) binding to the *Nme*Cas9 HNH surface to the corresponding, engineered residues (15Q (left), 3F and 48I (right)) binding to the *Sau*Cas9 HNH surface.

Next, we characterized AcrX performance in detail by targeting *Sau*Cas9 to different loci (Fig. 4a). AcrX efficiently suppressed *Sau*Cas9 genome editing at all tested loci, showing up to a ∼30-fold improvement in inhibition as compared to the parental AcrIIC1 (Fig. 4b, Supplementary Figs. 13 and 14). Interestingly, akin to wild-type AcrIIC1, mCherry domain fusion to AcrX loop 5 strongly improved inhibition on *Nme*Cas9 (Supplementary Fig. 12), but had no noticeable effect on *Sau*Cas9 inhibition. This indicates that the potent inhibition mediated by our chimeric Acrs arises from specific features of the *Nme*Cas9 structural architecture.

**Fig. 4.**
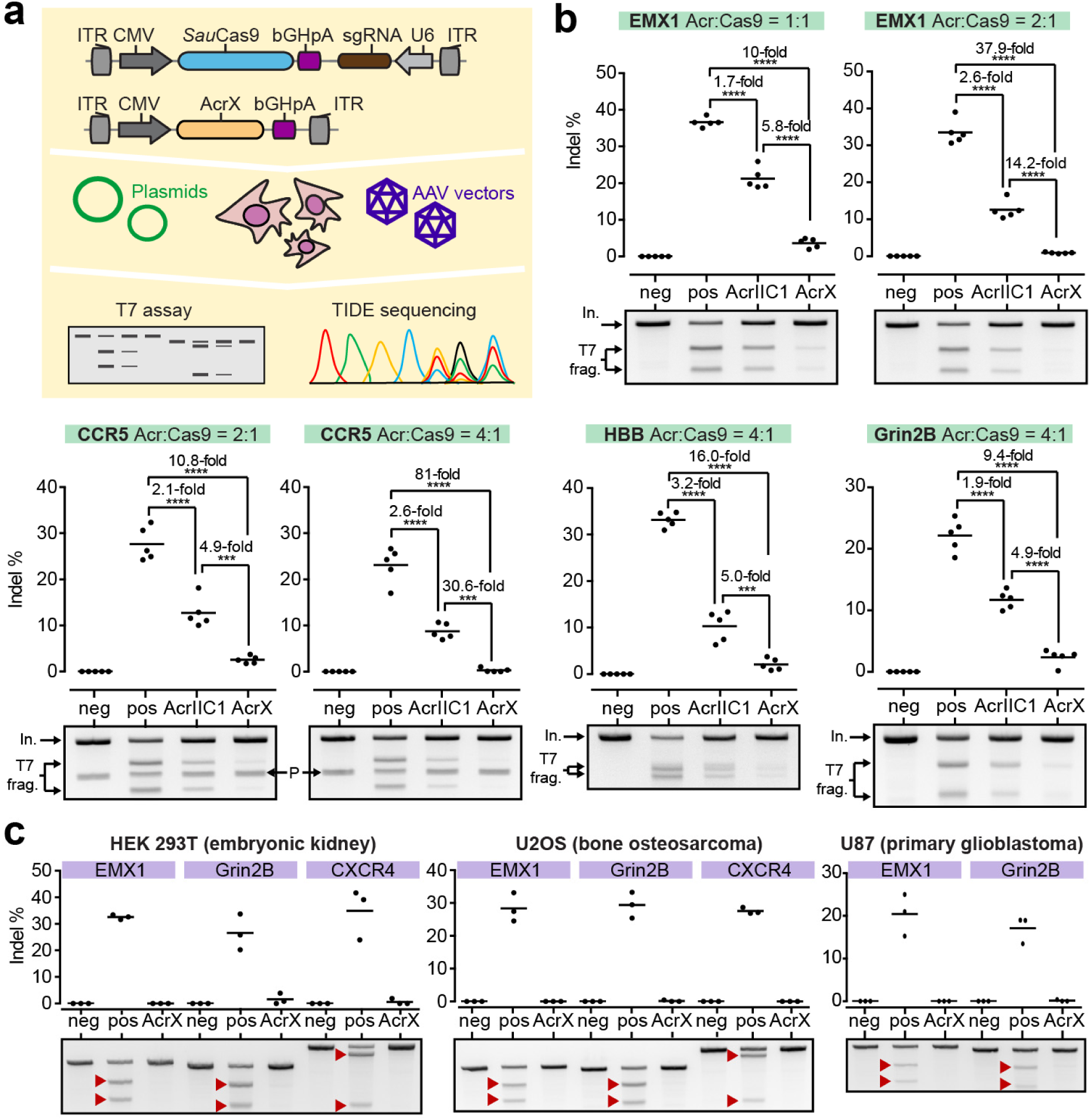
Characterization of AcrX, an engineered *S. aureus* Cas9 inhibitor. (**a**) Schematics of vectors and experimental setup. (**b**) HEK 293T cells were co-transfected with the vectors in **a** targeting the indicated loci followed by T7 endonuclease assay. The Acr:Cas9 vector ratio used during transfection is indicated. P denotes a T7 cleavage band which is due to a polymorphism in the CCR5 gene (Supplementary Fig. 15). In., input band. T7 frag., T7 cleavage fragments. \*\*\**P* < *0.001*, \*\*\*\**P < 0.0001* by one-way ANOVA with Bonferroni correction. (**c**) AAV-mediated delivery of AcrX results in potent *Sau*Cas9 inhibition in different cell lines. Cells were co-transduced with AAV2 vectors expressing (i) *Sau*Cas9 and a sgRNA targeting the indicated loci and (ii) AcrX followed by T7 endonuclease assay. Red triangles point to T7 cleavage fragments. (**b**,**c**) Representative T7 gel images and corresponding quantification of indel frequencies are shown. Lines in the plots indicate means, dots individual data points for n = 5 (**b**) or n = 3 (**c**) independent experiments. Neg, negative control (Cas9 only). Pos, positive control (Cas9 + sgRNA).

Finally, to test the performance of AcrX in different mammalian cell lines and upon viral delivery, we packaged (i) AcrX as well as (ii) *Sau*Cas9 and sgRNAs targeting the EMX1, Grin2B or CXCR4 locus into AAV serotype 2. We then transduced HEK 293T (human embryonic kidney), U2OS (human osteosarcoma) or U87 (human primary glioblastoma) cells with these vectors and found that indel formation was reduced to undetectable levels in practically all samples that received AcrX (Fig. 4c).

Here, we reported the engineering of anti-CRISPR proteins with improved inhibition potency and expanded target specificity. We showed that domain insertion into AcrIIC1 enhances *Nme*Cas9 inhibition, likely by interfering with DNA binding. On top, using structure-based design, we created the first anti-CRISPR protein able to efficiently block *Sau*Cas9 function. Our findings show that structure-guided protein engineering can yield anti-CRISPR proteins with desired properties that go beyond the currently known natural repertoire. These designer anti-CRISPRs might not only find wide application in the context of biotechnology and CRISPR-based therapies ^7-10^, but also provide an innovative strategy to safeguard CRISPR technology.

## Acknowledgments

We thank the Synthetic Biology (IPMB, Heidelberg University), Virus-host interactions (Heidelberg University clinics) and the Protein Design and Immunoengineering groups (EPFL, Lausanne) for helpful discussions, Kathleen Börner (Heidelberg University clinics, Heidelberg, Germany) and Karsten Rippe (DKFZ, Heidelberg, Germany) for providing cell lines and Katharina Niopek for critical reading of the manuscript. We would like to thank the EPFL’s Scientific IT and Application Support Center for their support on computational infrastructure. We would like to thank the Protein Production and Structure Core facility for their support on the protein biophysical characterization experiments. This study was funded by the Helmholtz association, the German Research Foundation (DFG) and the Federal Ministry of Education and Research (BMBF) (R.E.). D.G. is grateful for funding from the German Center for Infection Research (DZIF, TTU-HIV 04.803 and TTU-HIV 04.815) and the Cystic Fibrosis Foundation Therapeutics (CFFT, grant number GRIMM15XX0). C.S., D.G. and R.E. acknowledge funding from the Transregional Collaborative Research Center TRR179 (DFG, Projektnummer 272983813). C.S. and D.G. acknowledge additional funding by the Cluster of Excellence CellNetworks (DFG, EXC81). B.E.C. is a grantee from the European Research Council (Starting grant - 716058), the Swiss National Science Foundation and the Biltema Foundation. The computational simulations were performed at the CSCS - Swiss National Supercomputing Centre through a grant obtained by B.E.C. Z.H. is supported by a grant from the Swiss National Science Foundation. S.R. is supported by a grant of the National Center of Competence in Research in Chemical Biology.

## Author contributions

D.N. conceived the initial idea and refined it together with J.U.z.B. and B.E.C.; J.M, C.S., S.A. and D.N. designed and performed experiments. S.A. and M.D.H. performed AAV production. S.R. purified the Acrs and performed protein-biochemical characterization. Z.H., J.U.z.B., A.S. and B.E.C. performed *in silico* structural analysis and modeling. D.G. provided expertise on AAVs and *Sau*Cas9. B.E.C. and D.N. jointly directed the work. B.E.C. and D.N. wrote the manuscript with support from all authors.

## Online Methods

### Modeling of AcrIIC1-mCherry fusions

We used the Rosetta remodel application^25^ to generate the AcrIIC1-mCherry chimeras based on the structures for AcrIIC1 (PDB 5VGB) and mCherry (PDB 4ZIN). The N- and C-termini of mCherry were absent in the crystal structure and were rebuilt using fragment insertion together with cyclic coordinate descent^26^ and kinematic closure^27, 28^ with default values. For the designed chimera with a two-residue deletion, approximately 1500 decoys were generated and subsequently clustered with a root mean square deviation (r.m.s.d) threshold of 5 Å into 27 clusters. For the chimera with additional GSG-linkers at the N- and C-termini, approximately 1200 structures were clustered in 100 clusters with the same parameters. Representative examples of the three most populated clusters, which also have the lowest energies, are shown to illustrate the potential structural diversity of the AcrIIC1-mCherry chimeras (Supplementary Fig. 3). Analysis of the Rosetta outputs, structural models and the biochemical data was performed using the rstoolbox^29^.

### AcrIIC1 interface design

To screen *in silico* for mutations that could enhance the affinity of AcrIIC1 to *Staphylococcus aureus* (*Sau*)Cas9, we modeled a complex of AcrIIC1 (originally crystalized with the HNH domain of *Neisseria meningitidis* (*Nme*)Cas9, PDB 5VGB) with the *Sau*Cas9 HNH domain (PDB 5CZZ). A first structural alignment was performed between the *Nme* and *Sau*Cas9 domains using TM-align^30^ revealing an r.m.s.d of 2.32 Å and several structurally and sequence-conserved interface regions. These conserved interface regions were then used to refine the alignment, using the PyMOL (version 2.3.1) superposition function, to obtain the modeled complex used for the design simulations (Supplementary Fig. 5). We then analyzed the interfaces of both orthologues to pinpoint hotspots which could be designed in AcrIIC1 to enhance its interaction with *Sau*Cas9. These hotspots were visualized with surface point-wise distances between the two surfaces that were computed with a custom script. For each point on the reference surface (*Nme*), the distance to the closest point on the other surface (*Sau*) was calculated. In the visualizations, these distances were binarized by setting a cutoff of 1 Å (Fig. 3).

Next, we used RosettaScripts^31^ to perform a single-site *in silico* mutagenesis, thereby allowing subsets of amino acids for each of the selected residues on AcrIIC1. From our interface analysis we selected the following residues in AcrIIC1 for in silico mutagenesis: N3, D15, R36, D43, D45, D46, K47, A48 and M77. The design protocol consisted of two rounds of packing and minimization with fixed backbone. We generated a total of 51 designs and computed their change in binding free energy (ddG), number of hydrogen bonds across the interface, change in hydrophobic solvent-accessible surface area (SASA), and interface shape complementary with the target *Sau*Cas9 HNH domain. Designs with improved ddGs compared to that of the AcrIIC1-*Sau*Cas9 complex (−22 Rosetta energy units), increased hydrogen bonds across the interface, improvements in SASA and shape complementarity were manually inspected. A total of 10 substitutions on 8 sites were selected for experimental validation. Mutations N3F, N3Y and A48I were designed to increase interface packing and pi-stacking with the complementary hydrophobic patches on *Sau*Cas9. D43F and D45F were generated to fill voids within the interface boundaries and increase the hydrophobic packing. D15Q, R36D, D46E, K47Q and M77S were introduced to balance the underlying charge distribution on *Sau*Cas9 within the respective region. The overall design workflow is shown in Supplementary Fig. 7.

After experimental validation and combination of the proposed mutations, the final AcrX (AcrIIC1 with N3F, D15Q, A48I) was modeled following the same protocol. Electrostatic properties for AcrIIC1, AcrX as well as the *Nme*Cas9 and the *Sau*Cas9 HNH domains were computed using the adaptive Poisson Boltzmann solver (APBS) plugin in PyMOL (Supplementary Fig. 10). The mutation D15Q results in a less negative potential, to optimize the interaction with a patch of the interface in *Sau*Cas9 that has a lower positive potential as compared to *Nme*Cas9.

### Protein expression and purification

DNA sequences of the designs were purchased from Twist Bioscience. For bacterial expression, the DNA fragments were cloned via Gibson cloning^32^ into a pET21b vector encoding a peptide sequence containing a tobacco etch virus (TEV) protease cleavage site followed by a terminal His-tag and transformed into *E. coli* BL21(DE3). Expression was conducted in Terrific Broth supplemented with ampicillin (100 μg per ml). Cultures were inoculated at an OD600 of 0.1 from an overnight culture and incubated in a shaker at 37°C and 220 rpm. After reaching an OD600 of 0.6, expression was induced by the addition of 0.4 mM IPTG and cells were further incubated overnight at 20°C. Cells were harvested by centrifugation and pellets were resuspended in lysis buffer (50 mM TRIS, pH 7.5, 500 mM NaCl, 5% glycerol, 1 mg per ml lysozyme, 1 mM PMSF, 4 μg per ml DNase). Resuspended cells were sonicated and clarified by centrifugation. Ni-NTA purification of sterile-filtered (0.22 μm) supernatant was performed using a 5 ml His-Trap FF column on an ÄKTA pure system (GE Healthcare). Bound proteins were eluted using an imidazole concentration of 300 mM. Concentrated proteins were further purified by size exclusion chromatography on a Hiload 16/600 Superdex 75 pg column (GE Healthcare) using PBS buffer (pH 7.4) as mobile phase.

### Circular dichroism (CD)

Far-UV circular dichroism spectra of AcrIIC1 and AcrX were collected between a wavelength of 190 nm to 250 nm on a Jasco J-815 CD spectrometer in a 1 mm path-length quartz cuvette. Proteins were dissolved in 10 mM phosphate buffer at concentrations between 20 μM and 40 μM. Wavelength spectra were averaged from two scans with a scanning speed of 20 nm per min and a response time of 0.125 sec. The thermal denaturation curves were collected by measuring the change in ellipticity at 220 nm from 20 to 90°C with 2 or 5°C increments.

### Size-exclusion chromatography combined with Multi-Angle Light-Scattering (SEC-MALS)

Multi-angle light scattering was used to assess the monodispersity and molecular weight of the proteins. Samples containing 50–100 μg of protein in PBS buffer (pH 7.4) were injected into a Superdex 75 10/300 GL column (GE Healthcare) using an HPLC system (Ultimate 3000, Thermo Scientific) at a flow rate of 0.5 ml per min coupled in-line to a multi-angle light scattering device (miniDAWN TREOS, Wyatt). Static light-scattering signal was recorded from three different scattering angles. The scatter data were analyzed by ASTRA software (version 6.1, Wyatt).

### Construct design and cloning

Constructs used in this study are listed in Supplementary Table 1. Sequences for all plasmids created in this study are provided as GenBank files in Supplementary Data. The following constructs were generated via classical restriction enzyme cloning or Golden Gate assembly^33^. Oligonucleotides and synthetic double-stranded DNAs were obtained from IDT. PCRs were performed either with Q5 Hot Start high-fidelity DNA polymerase (New England Biolabs) or Phusion Flash high-fidelity polymerase (Thermo Fisher Scientific). After separating PCR products or restriction digest products on agarose gels, bands of the desired size were cut out and the DNA was extracted using the QIAquick gel extraction kit (Qiagen). Restriction enzymes and T4 DNA ligase were obtained from Thermo Fisher Scientific. Constructs were transformed into chemical competent Top10 cells (Thermo Fisher Scientific). DNA was purified using the QIAamp DNA Mini, Plasmid Plus Midi or Plasmid Maxi kit (all from Qiagen). The plasmid pEJS654 All-in-One AAV-sgRNA-hNmeCas9 co-encoding *Nme*Cas9 and a corresponding sgRNA expression cassette was a kind gift from Erik Sontheimer (Addgene #112139). The plasmid pX601-AAV-CMV::NLS-SaCas9-NLS-3xHA-bGHpA;U6::BsaI-sgRNA co-encoding *Sau*Cas9 and a corresponding sgRNA expression cassette was a kind gift from Feng Zhang (Addgene #61591). The luciferase reporter plasmid was previously reported by us^8^ and modified as follows: an NTS33 target site^34^ was inserted behind the firefly luciferase start codon and in frame with the luciferase gene; an NTS33-targeting sgRNA was subsequently inserted into the modified reporter. Vectors encoding AcrIIC1, AcrIIC3 and AcrIIA4 were previously reported by us^8, 9^. *As*LOV2-, PDZ- and mCherry-encoding sequences were obtained from IDT as human-codon-optimized synthetic DNA fragments (gBlocks). AcrIIC1 chimeras were created by Golden Gate cloning as follows: the plasmid encoding AcrIIC1 was first linearized at a selected position in the AcrIIC1 coding sequence via around-the-horn PCR; LOV2-, PDZ- and mCherry-coding sequences were then amplified by matching primers introducing optional GS linker encoding sequences and ligated into the linearized vector backbone; point mutations and protein tags were introduced via around-the-horn PCR via the primer overhangs. Annealed oligonucleotides corresponding to the target site sequence were cloned into the hybrid Cas9-sgRNA vectors via SapI (*Nme*Cas9) or BsaI (*Sau*Cas9) restriction sites as described previously^20, 34^.

### Cell culture and AAV lysate production

HEK 293T (human embryonic kidney), U87 (human primary glioblastoma; kindly provided by Kathleen Börner, Heidelberg University clinics) and U2OS (human osteosarcoma; kindly provided by Karsten Rippe, German Cancer Research Center (DKFZ), Heidelberg) were cultured at 5% CO_2_ and 37°C in a humidified incubator and maintained in phenol red-free Dulbecco’s Modified Eagle Medium (DMEM; ThermoFisher/GIBCO) supplemented with 10% (v/v) fetal calf serum (Biochrom AG), 2 mM L-glutamine and 100 U per ml penicillin and 100 µg per ml streptomycin (both ThermoFisher/GIBCO). The U2OS medium was additionally supplemented with 1 mM sodium pyruvate (GIBCO). Cell lines were free of mycoplasma contamination and authenticated prior usage (Multiplexion, Heidelberg, Germany).

AAV-containing cell lysates were produced by seeding HEK 293T cells into 6-well plates (Corning) at a density of 350,000 cells per well. On the next day, cells were co-transfected using 8 µl Turbofect reagent (ThermoFisher Scientific) per well and 1,333 ng of each of the following plasmids: (i) An AAV vector plasmid carrying the transgenes to be delivered flanked by inverted terminal repeats (ITRs), (ii) an AAV helper plasmid carrying *rep* and *cap* genes of AAV serotype 2 and (iii) an adenoviral helper plasmid providing the required helper functions^35^. The AAV vector plasmid encoded either (1) a dual transgene cassette expressing *Sau*Cas9 driven from a CMV promoter and sgRNA driven from a shortened U6 promoter, targeting either the EMX1, Grin2B or CXCR4 locus, in a single-stranded AAV context, (2) the same *Sau*Cas9 cassette but together with an empty sgRNA expression cassette (negative control) or (3) a CMV promoter-driven AcrX transgene in a double-stranded AAV context. Three days after transfection, cells were harvested in 300 µl PBS and lysed by subjecting them to five alternating freeze-thaw cycles in liquid nitrogen and in a 37°C water bath. Cell debris was separated by centrifugation and the supernatant containing the AAVs was stored at 4°C (for a maximum of two weeks) prior to usage.

### Luciferase reporter assays

HEK 293T cells were seeded into 96-well plates at a density of 12,500 cells per well. For titration experiments employing the chimeric Acrs (Fig. 2c), the cells were co-transfected on the following day with (i) 33 ng of a dual luciferase reporter plasmid encoding a firefly and *Renilla* luciferase gene and an sgRNA targeting the NTS33 site in the firefly reporter gene, (ii) 33 ng of a vector co-expressing *Nme*Cas9 and a sgRNA targeting the NTS33 site, (iii) 99, 33, 16.5, 6.6, 3.3 or 1.65 ng of Acr vector, and (iv) 0, 66, 82.5, 92.4, 95.7 or 97.35 ng of an irrelevant stuffer plasmid (pBluescript), respectively. The stuffer plasmid was added to keep the total amount of DNA transfected constant in all samples. For d*Nme*Cas9 or n*Nme*Cas9 experiments (Supplementary Fig. 4), 33 ng of the dual luciferase reporter, 33 ng of d- or n*Nme*Cas9 and 33 ng Acr construct were co-transfected. Transfections were performed using Lipofectamine 3000 reagent (Thermo Fisher Scientific) according to the manufacturer’s protocol.

Two days post-transfection, the cells were washed with 1x PBS and lysed with 30 µl passive lysis buffer (Promega) for 30 min, while being shaken on a thermomixer (Eppendorf) at 500 rpm and at room temperature. Finally, luciferase activity was analyzed using the Dual-Glo luciferase assay system (Promega). In short, 10 µl of lysate were transferred to a white sample plate and photo counts were measured with a GLOMAX 96 microplate luminometer (Promega). Integration time was 10 s with a delay of 2 s between substrate injection and measurement. To calculate the reported luciferase activity values, firefly luciferase photon counts were normalized those obtained for *Renilla* luciferase.

### T7 endonuclease assay and TIDE sequencing

Genomic target sites relevant for T7 and TIDE experiments are listed in Supplementary Table 2. For transfection-based experiments, HEK 293T cells were seeded in 96-well plates (Eppendorf) at a density of 12,500 cells per well. For AAV transduction-based experiments, HEK 293T, U2OS and U87 cells were seeded at a density of 3,500, 3,000 and 3,000 cells per well, respectively. For experiments with *Sau*Cas9 (Fig 4b, Supplementary Figs. 13 and 14), transfections were performed with JetPrime using 0.3 µl of JetPrime reagent per well, except for the experiment shown in Supplementary Fig. 12, in which Lipofectamine 3000 was employed for transfection of all samples including those with *Sau*Cas9. Note that the CXCR4 target site in Supplementary Fig. 13 is the CXCR4-1 site in Supplementary Table 2. For *Nme*Cas9 experiments (Fig. 2a,b), transfections were conducted with Lipofectamine 3000 using 0.2 µl Lipofectamine reagent, 0.4 µl p3000 and 200 ng total DNA per well. Cells were co-transfected with 100, 133 or 160 ng of Acr vector and 100, 67 or 40 ng or all-in-one Cas9/sgRNA vector corresponding to Acr:Cas9 vector ratios of 1:1, 2:1 and 4:1, respectively, as indicated in the figures. Transfections for the initial screen of the chimeric AcrIIC1 variants (Supplementary Fig. 2) were performed with only 100 ng of total DNA per well, using a 1:1 ratio of Cas9/sgRNA and Acr vectors.

For AAV-based experiments (Fig. 4c), cells were co-transduced with 50 µl of AcrX and 50 µl Cas9/sgRNA AAV lysates on two subsequent days. As negative and positive controls, cells were transduced with 50 µl of Cas9-only AAV lysate or Cas9/sgRNA AAV lysate (see above), respectively, topped up to 100 µl with PBS (to keep the transduction volume identical in all samples). Note, the CXCR4-target site in Fig. 4c is the CXCR4-2 site in Supplementary Table 2. Three days post-transfection or (initial) transduction, cells were harvested in DirectPCR Lysis Reagent (Peqlab) supplemented with Proteinase K (Sigma), and incubated at 55°C for at least 6 hours followed by Proteinase K inactivation at 85°C for 45 min. The CRISPR-Cas9-targeted genomic loci were then amplified via PCR with appropriate primers (Supplementary Table 3) using Q5 Hot Start High-Fidelity DNA Polymerase (New England Biolabs). Indel frequencies were assessed by T7 endonuclease assay or TIDE sequencing^36^.

For T7 assays, 5 µl of the target amplicons were diluted 1:4 in 1x NEB buffer 2 and subsequently denatured at 95°C for 5 min and re-annealed by applying a ramp rate of -2°C per sec at 95 to 85°C and -0.1°C per sec at 85 to 25°C using a nexus GSX1 Mastercycler (Eppendorf). Subsequently, 0.5 µl of T7 endonuclease (New England Biolabs) was added, and samples were incubated at 37°C for 15 min, followed by analysis on a 2% TBE agarose gel. The PCR input and T7 cleavage fragment bands were then quantified using the gel analysis tool in ImageJ^37, 38^ (http://imagej.nih.gov/ij/). The frequency of insertions and deletions was calculated using the formula indel(%) = 100 × (1 – (1-fraction cleaved)^1/2^), whereas the fraction cleaved = Sum(cleavage product bands)/Sum(cleavage product bands + PCR input band). Full-length gel images are shown in Supplementary Fig. 16.

For TIDE sequencing analysis, the target locus PCR amplicon was purified from a 1% agarose gel using the QIAquick Gel Extraction Kit (Qiagen). The DNA concentration was determined using a nano-photometer (Nanodrop, Thermo Fisher Scientific) and DNA was diluted to a final concentration of 75 ng per µl and sent for Sanger sequencing (Eurofins, Germany). Percentages of modified sequences were then quantified using the TIDE web tool (https://tide.deskgen.com/).

### Statistical analysis

Individual data points correspond to independent experiments with cells that were seeded and transfected/transduced independently and on different days. Each data point shown for the luciferase experiments further represents the mean of three technical replicates, *i.e*. cell cultures in different wells that were transfected and treated in parallel. Reported differences between groups were analyzed for statistical significance by one-way ANOVA and Bonferroni’s corrected post-hoc test. One, two, three and four asterisks thereby indicated *P* values below 0.05, 0.01, 0.001 and 0.0001, respectively. *P* values < 0.05 were considered statistically significant.

### Reporting Summary

Further information on experimental design is available in the Life Sciences Reporting Summary linked to this article.

### Data and code availability

Vectors encoding AcrX, AcrX* as well as the AcrIIC1-mCherry chimera will be made available via Addgene (plasmids # 128112-128114). Annotated vector sequences (GenBank files) are provided in Supplementary Data. Code and data for the computational domain assembly and the design of the improved AcrIIC1 point mutants will be made available on GitHub (https://github.com/zanderharteveld/AcrX). Additional data can be obtained from the corresponding authors on request.

