## Supplementary Information for "Computational design of anti-CRISPR proteins with improved inhibition potency and expanded specificity"

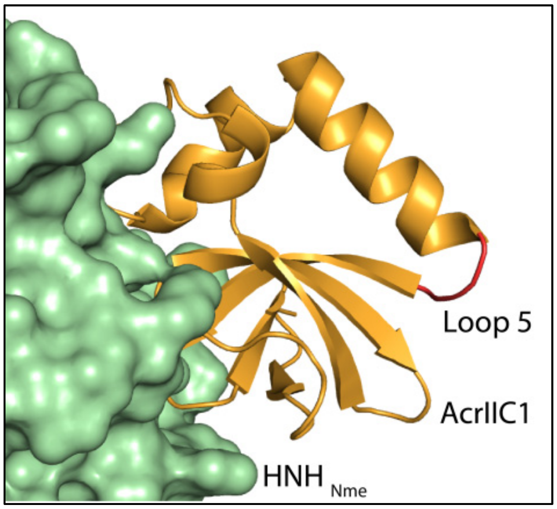


**Supplementary Figure 1 | Identification of a loop amenable to domain insertion.**
Loop 5 (in red) is located on the opposite site of the HNH-domain binding interface. It connects an alpha-helix and a beta-sheet via two residues (Tyrosine 70, Alanine 71). Structures shown correspond to PDB 5VGB.

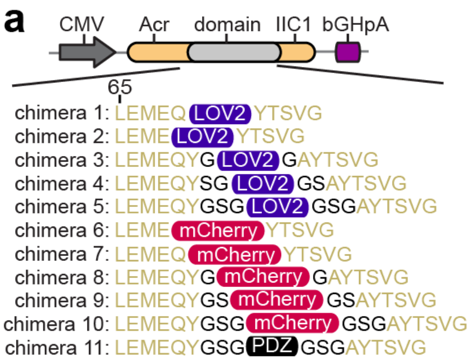


**Supplementary Figure 2 | Screening of AcrIIC1 domain insertion variants.**

(**a**) Schematic of chimeric Acrs. The chimeras comprise AcrIIC1 bearing an *Avena sativa* LOV2, mCherry or PDZ domain inserted into loop 5. The constructs carry optional GS linkers flanking the inserted domain (black residues) or deletions in loop 5. AcrIIC1 residue L65 is indicated. (**b**) Screen of chimeric Acrs in HEK 293T cells. Cells were co-transfected with vectors expressing *Nme*Cas9, the indicated Acr and a sgRNA targeting the IL2RG locus followed by T7 endonuclease assay. The Acr:Cas9 vector ratio during transfection was 1:1. Bars indicated means, error bars the SD and dots individual data points for n = 3 independent experiments. Numbers correspond to the constructs shown in **A.** Neg, negative control (Cas9 only). Pos, positive control (Cas9 + sgRNA).

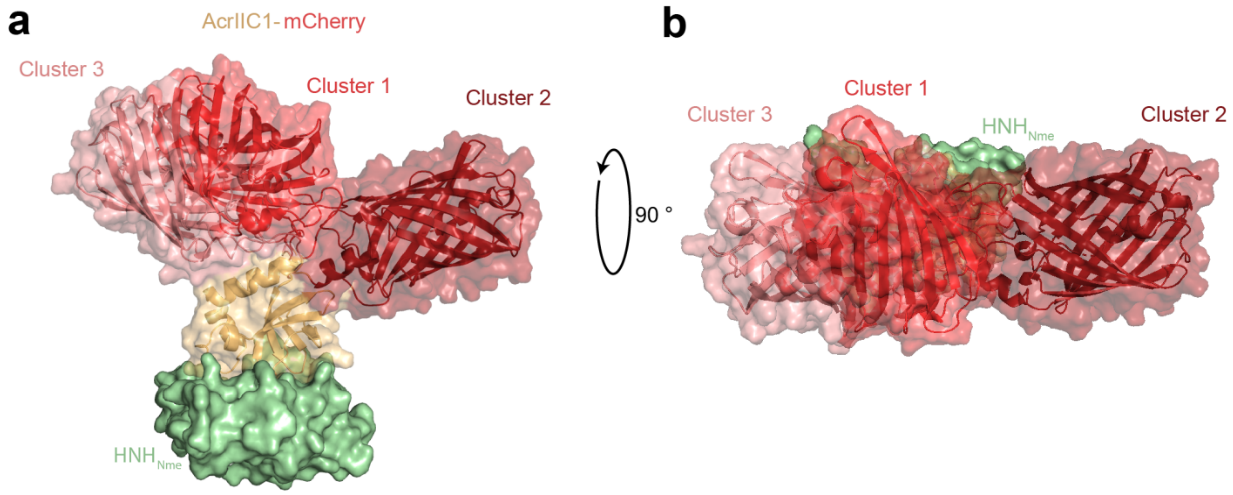


**Supplementary Figure 3 | Acr-mCherry chimera conformations.**
For the shown chimera, the mCherry was inserted after residue Q69 and before Y72. Residues Y70 and A71 were deleted. Representative examples of the three most populated clusters are shown, aligned to the structure of AcrIIC1 in complex with the *Nme*Cas9 HNH domain. (**a**) Side view. (**b**) Top view. The models are based on PDB 5VGB and 4ZIN.

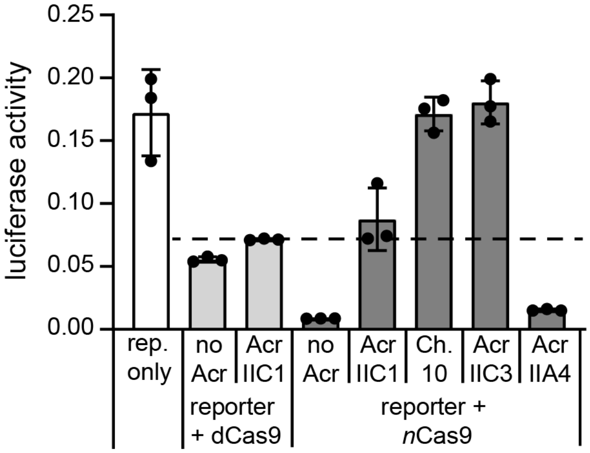


**Supplementary Figure 4 | The AcrIIC1-mCherry chimera, but not wild-type AcrIIC1 impairs *Nme*Cas9 DNA binding.**

HEK 293T cells were co-transfected with constructs expressing (i) a firefly luciferase reporter and sgRNA targeting the firefly luciferase gene, (ii) d*Nme*Cas9 or *Nme*Cas9 nickase (n*Nme*Cas9) and (iii) the indicated Acr followed by luciferase assay. The dotted line indicates the level of reporter knockdown due to CRISPRi by d*Nme*Cas9 (in presence of AcrIIC1). Bars indicate means, error bars the SD for n = 3 independent experiments. Rep. only, firefly luciferase reporter only control. See Supplementary Text for details.

```
Sau    1 REKNSKDAQKMINEMQKRNRQTNERIEEIIRT-----TGKENAKYLIEKI 45
            :.|| :|.|.:.|:.||:..|:.....|.     .|:..:|.:: |:
Nme    1 ---SFKD-RKEIEKRQEENRKDREKAAAKFREYFPNFVGEPKSKDIL-KL 45

Sau   46 KLHDMQEGKCLYSLEAIPLEDLLNNPFNYEVDHIIPRSVSFDNSFNNKVL 95
         :|::.|.||||||.:.|.| ..||.....|:||.:|.|.::|:|||||||
Nme   46 RLYEQQHGKCLYSGKEINL-GRLNEKGYVEIDHALPFSRTWDDSFNNKVL 94

Sau   96 VKQEENSKKGNRTPFQYLSSSDSKISYETFKKHILNLAKGKGRISKTKKE 145
         |...||..|||:||::|.:..|:...::.||..:     ...|..::||:
Nme   95 VLGSENQNKGNQTPYEYFNGKDNSREWQEFKARV-----ETSRFPRSKKQ 139

Sau  146 YLLEERDINRFSVQKDFINRNL 167
         .:|    :.:|. :..|..|||
Nme  140 RIL----LQKFD-EDGFKERNL 156
```

**Supplementary Figure 5 | Alignment of the *Sau*- and *Nme*Cas9 HNH domains.**
Conserved residues within the AcrIIC1-binding interface are in red. Bold characters indicate residues that were used to align the interfaces.

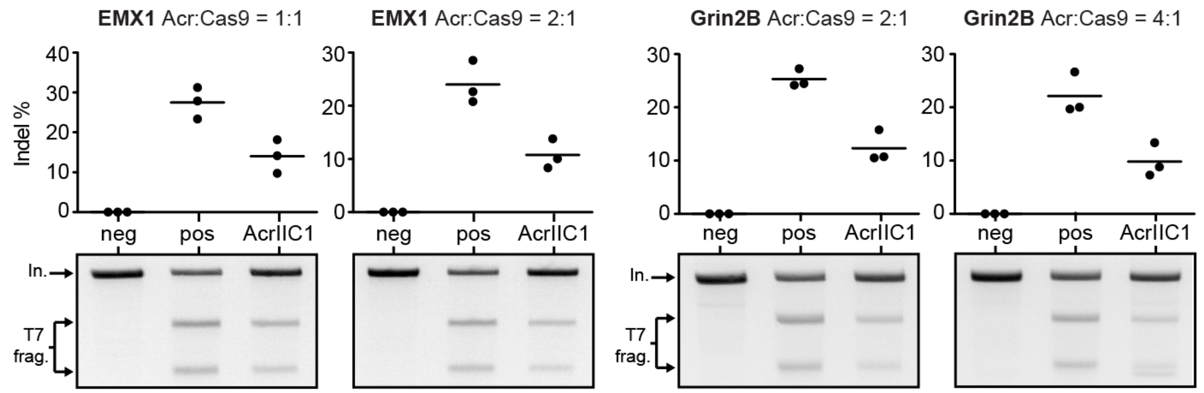


**Supplementary Figure 6 | AcrIIC1 is able to partially inhibit genome editing by *S. aureus* Cas9.**

HEK 293T cells were co-transfected with the vectors expressing *Sau*Cas9, a sgRNA targeting the EMX1 or Grin2B locus and AcrIIC1 followed by T7 endonuclease assay. The Acr:Cas9 vector ratio used during transfection is indicated. Lines in the plots indicate means, dots individual data points for n = 3 independent experiments. In., input band. T7 frag., T7 cleavage fragments. Neg, negative control (Cas9 only). Pos, positive control (Cas9 + sgRNA).

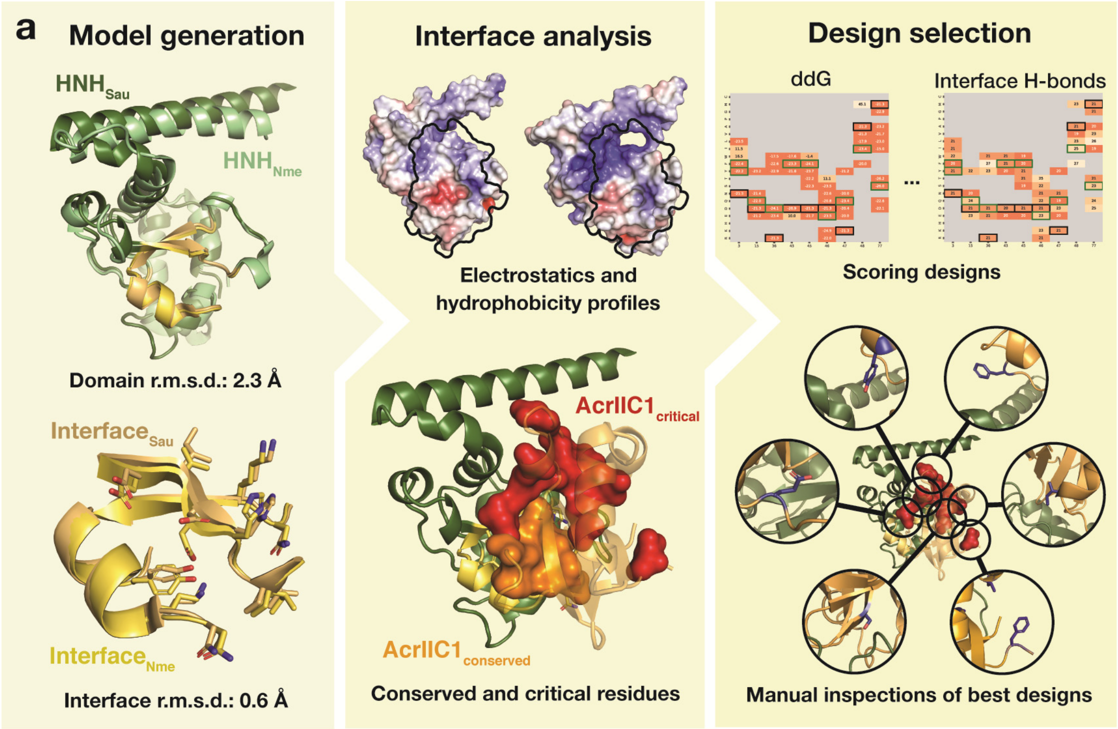


**b**

```
AcrIIC1    1   MANKTYKIGKNAGYDGCGLCLAAISENEAIKVKYLRDIC
AcrX       1   MAFKTYKIGKNAGYQGCGLCLAAISENEAIKVKYLRDIC
                 3            15
AcrIIC1   40   PDYDGDDKAEDWLRWGTDSRVKAAALEMEQYAYTSVGMA
AcrX      40   PDYDGDDKIEDWLRWGTDSRVKAAALEMEQYAYTSVGMA
                       48
AcrIIC1   79   SCWEFVEL
AcrX      79   SCWEFVEL
```

**Supplementary Figure 7 | Computational interface design.**

(**a**) After generating the *Sau*Cas9 HNH – AcrIIC1 model by structural alignment of the *Nme*Cas9 and *Sau*Cas9 HNH domains and analysis of the interfaces, conserved and critical residues to be kept or mutated were determined. Single-site *in silico* mutation experiments were performed and top scoring variants were experimentally validated after manual inspection. (**b**) Sequence alignment of AcrIIC1 and AcrX. Mutations are marked in bold red characters.

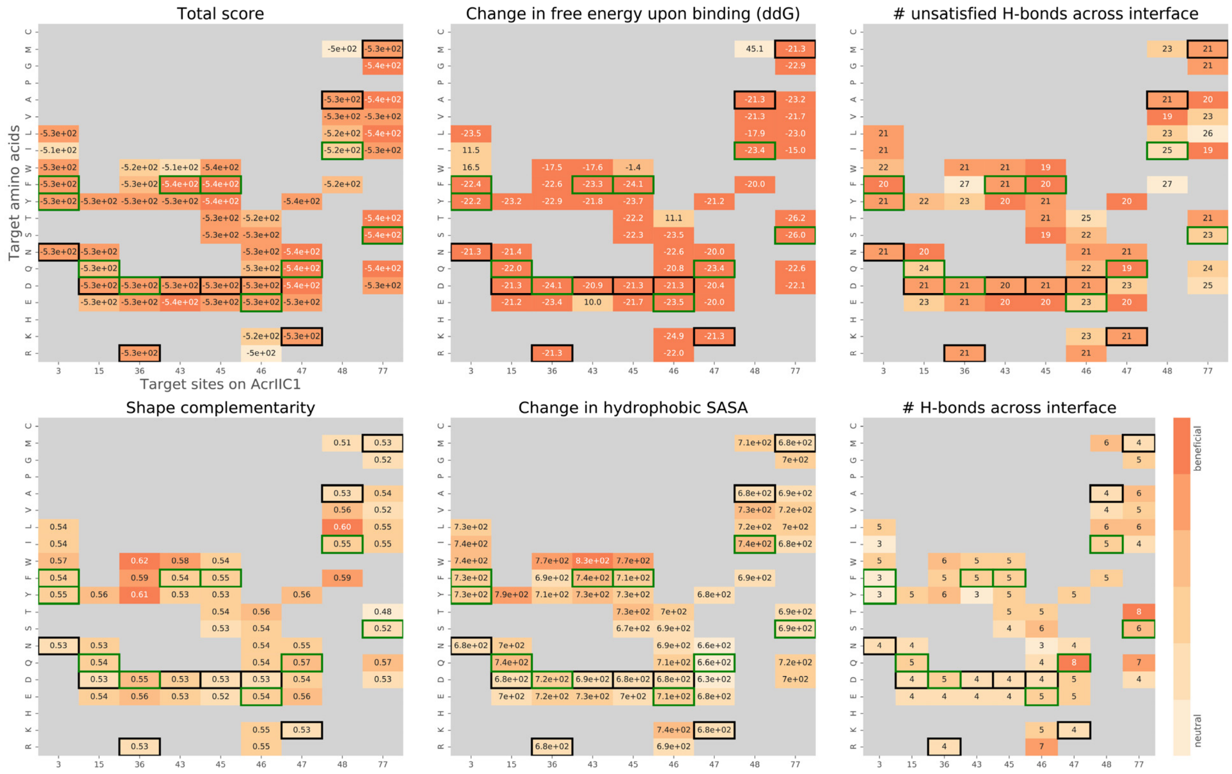


**Supplementary Figure 8 | Rosetta scores of single site mutants.**
Heatmaps of the computed structural metrics for the generated designs. Black boxes refer to the wild-type (AcrIIC1) scores per position, while green boxes indicate scores of selected designs for experimental validation.

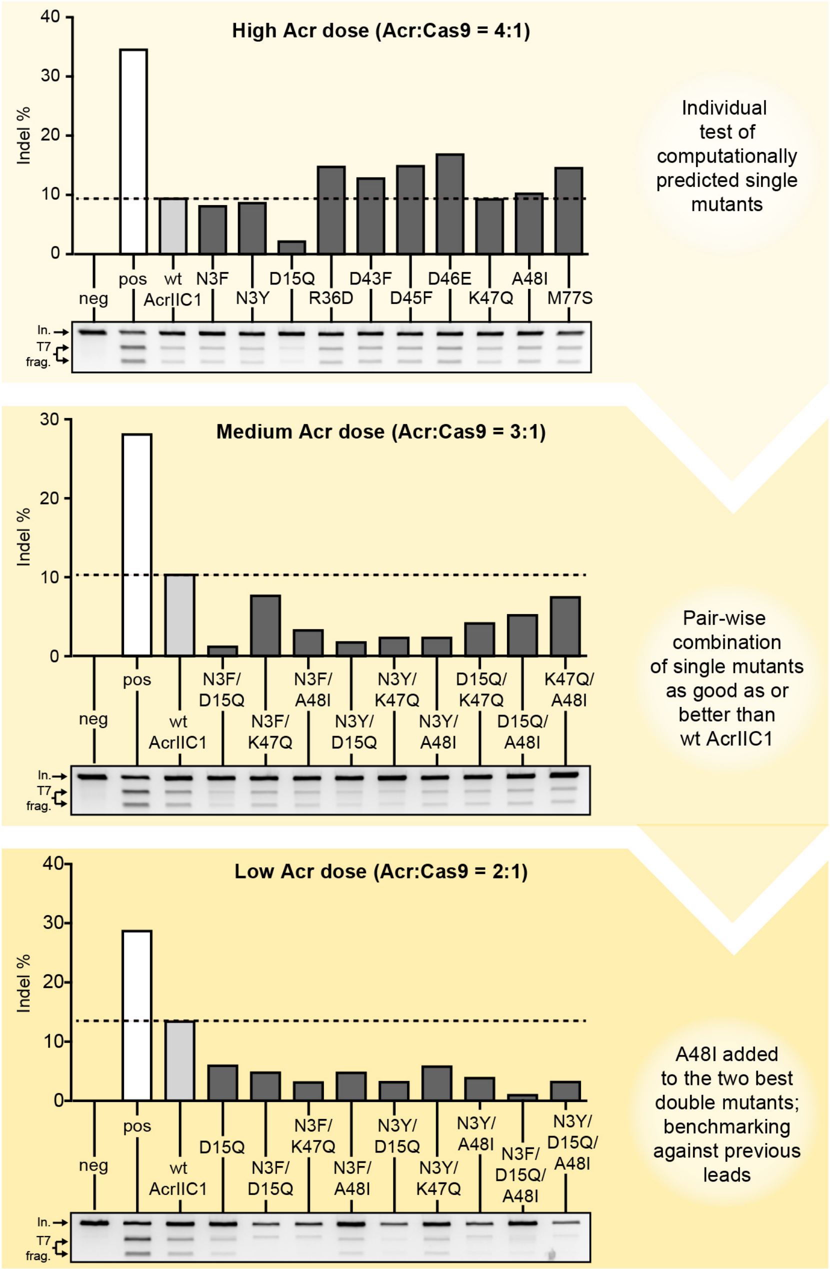


**Supplementary Figure 9 | Screening and iterative optimization of computationally designed AcrIIC1 mutants.**

HEK 293T cells were co-transfected with the vectors expressing *Sau*Cas9, a sgRNA targeting the EMX1 locus and either wild-type (wt) AcrIIC1 or the indicated AcrIIC1 mutant followed by T7 endonuclease assay. The Acr:Cas9 vector ratio (indicated) used during transfection was decreased with every iteration. Representative T7 gel images and corresponding quantifications of indel frequencies are shown. Dotted lines indicate the editing frequency in the presence of wild-type AcrIIC1. In., input band. T7 frag., T7 cleavage fragments. Neg, negative control (Cas9 only). Pos, positive control (Cas9 + sgRNA).

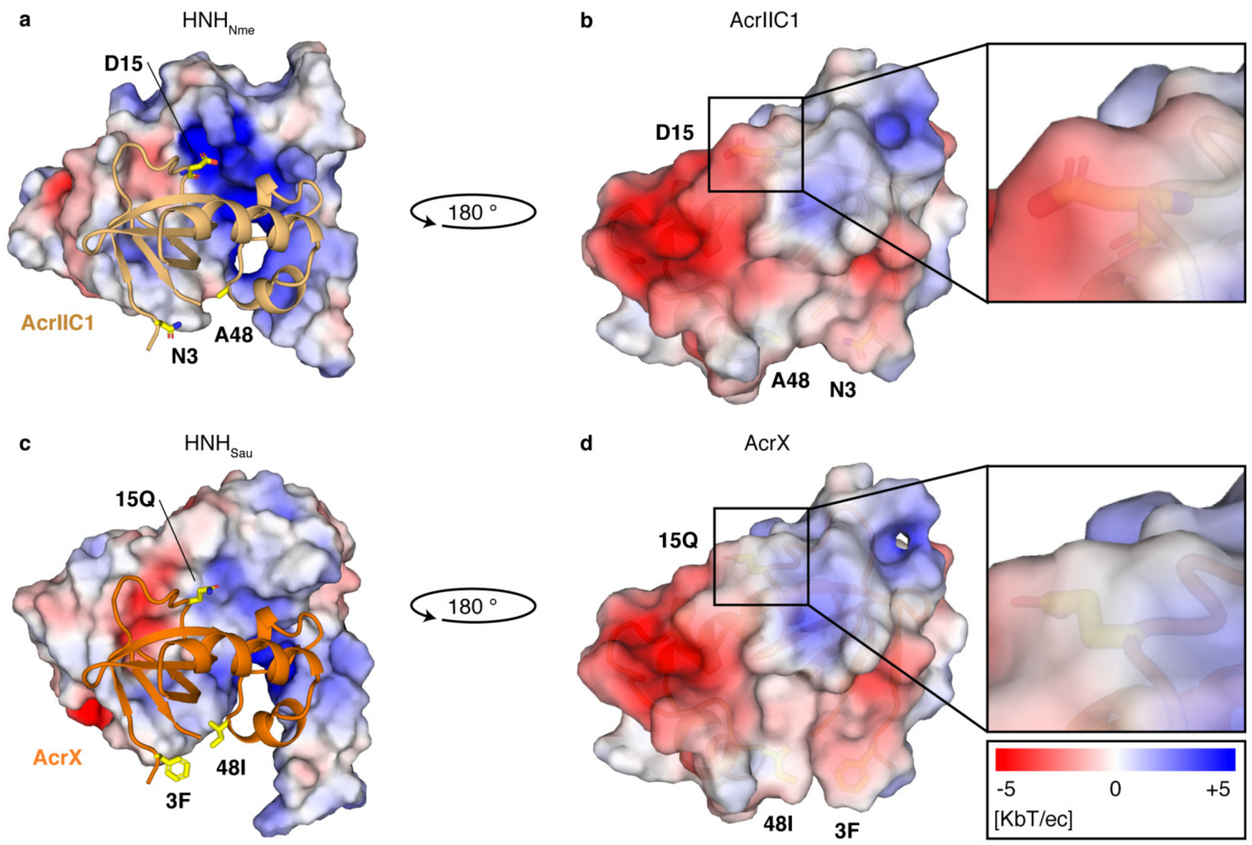


**Supplementary Figure 10 | Electrostatic potential of HNH domain and AcrIIC1 surfaces**. Potentials were calculated using the Adaptive Poisson-Boltzmann Solver. (**a**) The *Nme*Cas9 HNH domain has a strong positive potential close to residue 15 of AcrIIC1 in bound state. (**b**) AcrIIC1 has a similarly strong negative potential in the Cas-binding site. Residue 15 is at the border of this negative patch. (**c**) The *Sau*Cas9 HNH domain shows reduced positive potential around AcrIIC1 residue 15 as compared to *Nme*Cas9. (**d**) Mutation of residue 15 from aspartic acid to glutamine reduces the negative potential of AcrIIC1 in this position. (PDB 5VGB). (**c,d**) AcrX is the AcrIIC1 N3F, D15Q, A48I triple mutant.

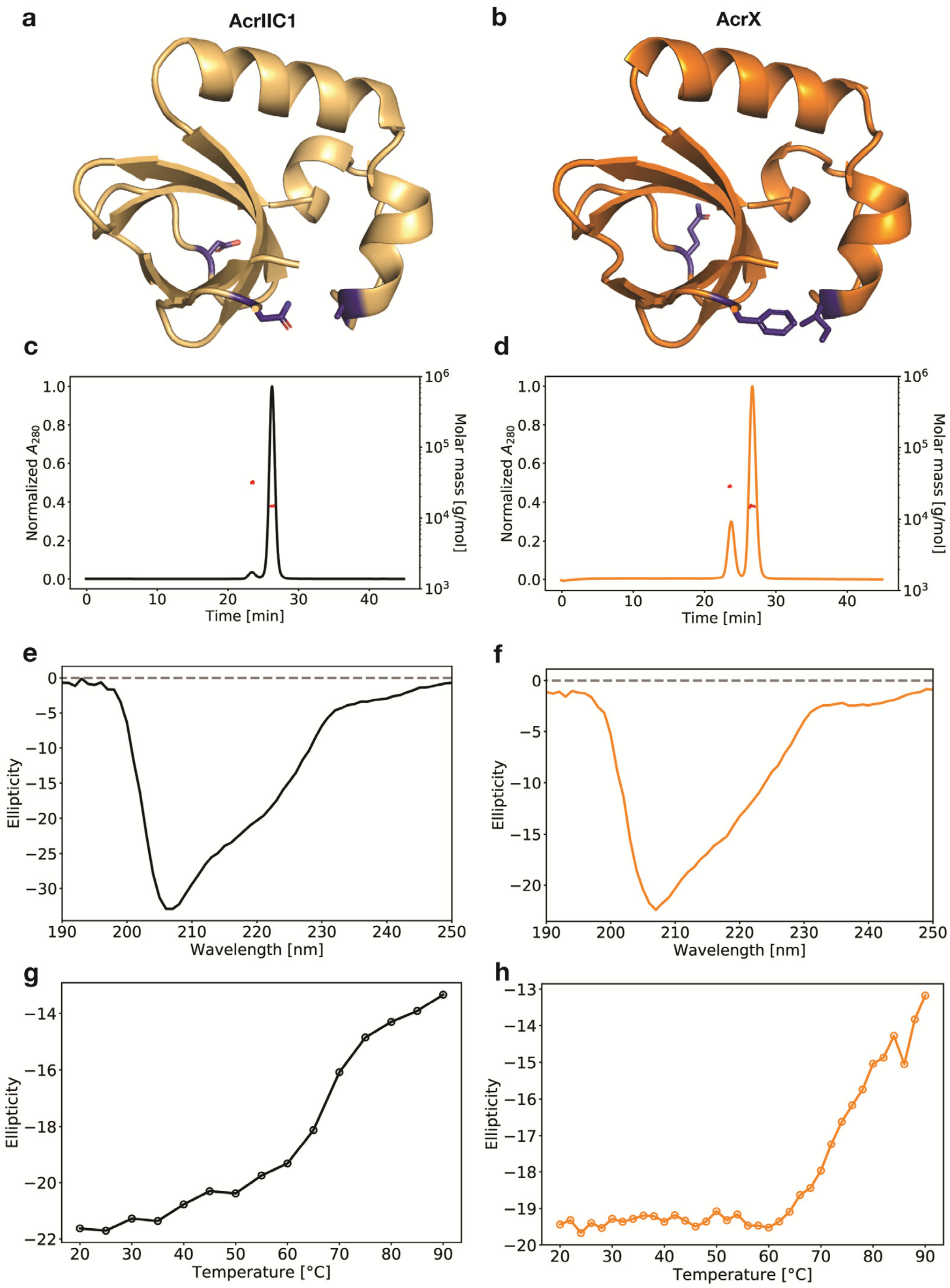


**Supplementary Figure 11 | Biochemical analysis of AcrIIC1 and AcrX.**
(**a**, **b**) Structure of AcrIIC1 and model of AcrX. Shown in blue are the three points mutations differentiating AcrIIC1 from AcrX. (**c**, **d**) SEC-MALS for AcrIIC1 and AcrX indicate a mainly monomeric form. AcrX shows a slightly higher number of dimers compared to AcrIIC1. (**e**, **f**) CD spectra for AcrIIC1 and AcrX show a peaked signal at around 207 nm as is typical for mixed alpha and beta secondary structures. (**g**, **h**) Thermal melting CD spectra for AcrIIC1 and AcrX are shown.

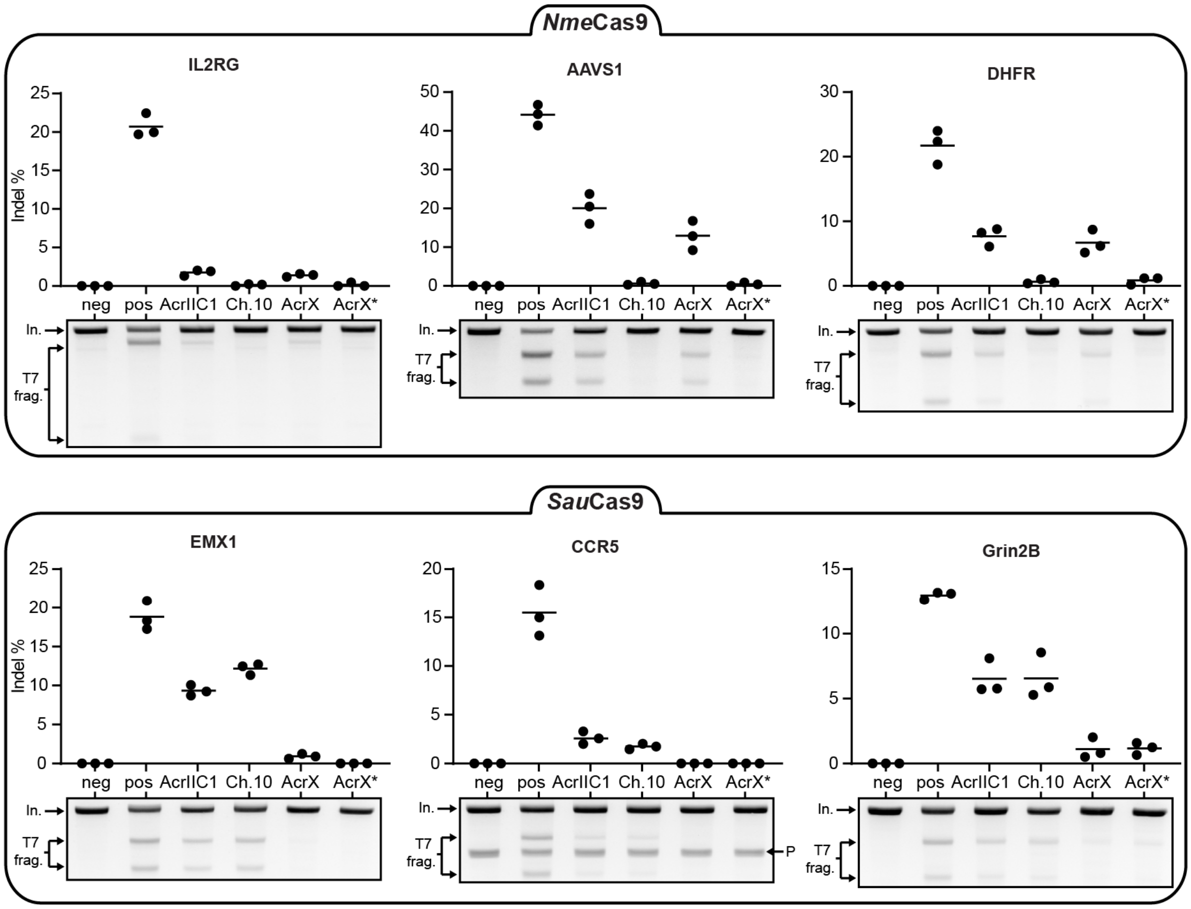


**Supplementary Figure 12 | AcrX inhibits *Nme*Cas9 function, which can be further improved by domain insertion.**

HEK 293T cells were co-transfected with vectors expressing (i) *Nme*Cas9 or *Sau*Cas9, (ii) a sgRNA targeting the indicated locus as well as the indicated Acr variant followed by T7 endonuclease assay. The Acr:Cas9 vector ratio used during transfection was 1:1 for the *Nme*Cas9 and 2:1 for the *Sau*Cas9 samples. Representative T7 gel images and corresponding quantification of indel frequencies are shown. Lines in the plots indicate means, dots individual data points for n = 3 independent experiments. In., input band. T7 frag., T7 cleavage fragments. Neg, negative control (Cas9 only). Pos, positive control (Cas9 + sgRNA). P denotes a T7 cleavage band which is due to a polymorphism in the CCR5 gene (Supplementary Figure 15). Ch. 10, Acr chimera #10 in Supplementary Figure 2. AcrX*, AcrX-mCherry chimera.

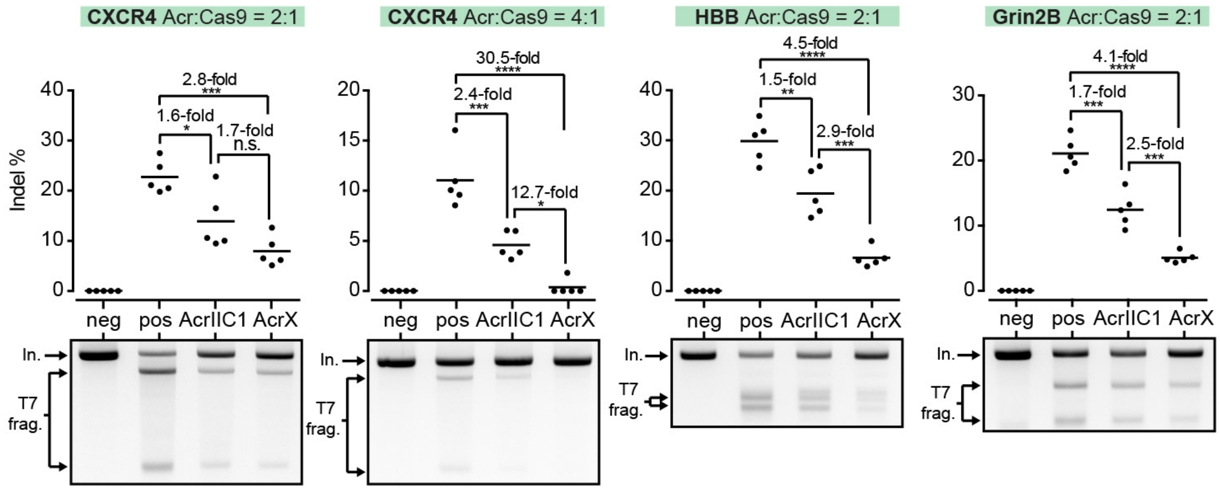


**Supplementary Figure 13 | AcrX is a potent *S. aureus* Cas9 inhibitor.**

HEK 293T cells were co-transfected with vectors expressing *Sau*Cas9, a sgRNA targeting the CXCR4, HBB or Grin2B locus and AcrIIC1 or AcrX followed by T7 endonuclease assay. The Acr:Cas9 vector ratio used during transfection is indicated. Representative T7 gel images and corresponding quantification of indel frequencies are shown. Lines in the plots indicate means, dots individual data points for n = 5 independent experiments. In., input band. T7 frag., T7 cleavage fragments. Neg, negative control (Cas9 only). Pos, positive control (Cas9 + sgRNA).

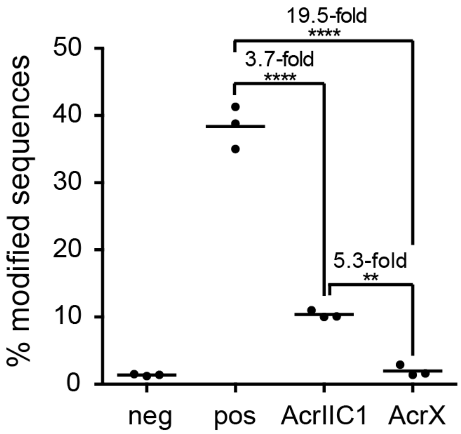


**Supplementary Figure 14 | TIDE sequencing analysis of *S. aureus* Cas9 inhibition by AcrX.** HEK 293T cells were co-transfected with plasmids expressing *Sau*Cas9, a sgRNA targeting the EMX1 locus and either AcrIIC1 or AcrX followed by TIDE sequencing[1]. Lines in the plots indicate means, dots individual data points for n = 3 independent experiments. Neg, negative control (Cas9 only). Pos, positive control (Cas9 + sgRNA). $^{**}P < 0.01$, $^{****}P < 0.0001$ by one-way ANOVA with Bonferroni correction.

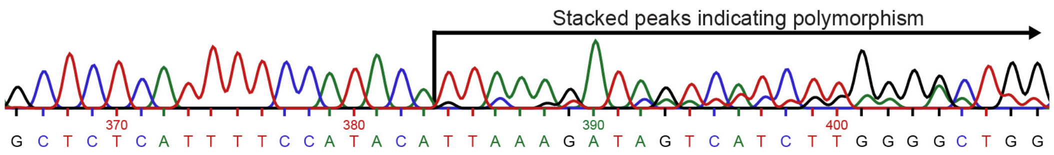


**Supplementary Figure 15 | Polymorphism in the CCR5 gene.**

Sanger sequencing confirms a polymorphism within the primer-flanked region in CCR5 (genomic DNA is derived from untreated HEK 293T cells).

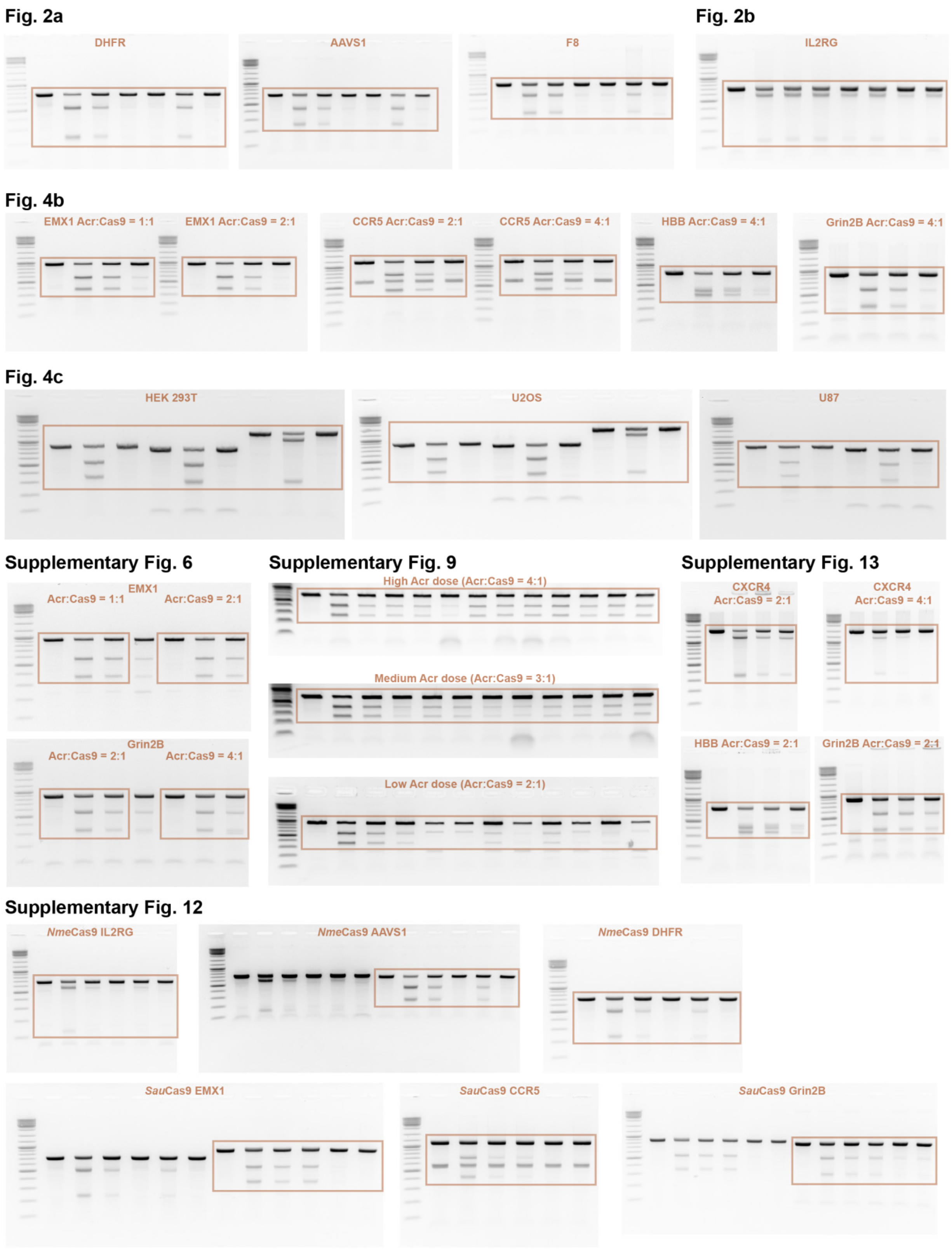


**Supplementary Figure 16 | T7 gel images**.
The ladder is the Gene Ruler DNA Ladder Mix (Thermo Fisher).

| # | Name | Insert | Source |
|---|---|---|---|
| 1 | pBluescript | empty vector | Invitrogen |
| 2 | luciferase reporter | firefly and *Renilla* luciferase + sgRNA targeting firefly gene | This work |
| 3 | hNmeCas9 + sgRNA scaffold | NLS hNmeCas9 NLS 3xHA; U6 promoter sgRNA scaffold | Reference [2] |
| 4 | hNmeCas9 + NTS33 sgRNA | NLS hNmeCas9 NLS 3xHA; U6 promoter NTS33 sgRNA | This work |
| 5 | dhNmeCas9 + NTS33 sgRNA | NLS catalytically dead hNmeCas9 NLS 3xHA; U6 promoter NTS33 sgRNA | This work |
| 6 | nhNmeCas9 + NTS33 sgRNA | NLS NmeCas9 nickase NLS 3xHA; U6 promoter NTS33 sgRNA | This work |
| 7 | hNmeCas9 + IL2RG sgRNA | NLS hNmeCas9 NLS 3xHA; U6 promoter IL2RG sgRNA | This work |
| 8 | hNmeCas9 + NTS4B sgRNA | NLS hNmeCas9 NLS 3xHA; U6 promoter NTS4B sgRNA | This work |
| 9 | hNmeCas9 + NTS59 sgRNA | NLS hNmeCas9 NLS 3xHA; U6 promoter NTS59 sgRNA | This work |
| 10 | hNmeCas9 + DHFR sgRNA | NLS hNmeCas9 NLS 3xHA; U6 promoter DHFR sgRNA | This work |
| 11 | hNmeCas9 + F8 sgRNA | NLS hNmeCas9 NLS 3xHA; U6 promoter F8 sgRNA | This work |
| 12 | hSaCas9 + sgRNA scaffold | NLS hSaCas9 NLS 3xHA; U6 promoter sgRNA scaffold | Reference [3] |
| 13 | hSaCas9 + EMX1 sgRNA | NLS hSaCas9 NLS 3xHA; U6 promoter EMX1 sgRNA | This work |
| 14 | hSaCas9 + CCR5 sgRNA | NLS hSaCas9 NLS 3xHA; U6 promoter CCR5 sgRNA | This work |
| 15 | hSaCas9 + GRIN2B sgRNA | NLS hSaCas9 NLS 3xHA; U6 promoter GRIN2B sgRNA | This work |
| 16 | hSaCas9 + HBB sgRNA | NLS hSaCas9 NLS 3xHA; U6 promoter HBB sgRNA | This work |
| 17 | hSaCas9 + CXCR4-1 sgRNA | NLS hSaCas9 NLS 3xHA; U6 promoter CXCR4-1 sgRNA | This work |
| 18 | hSaCas9 + CXCR4-2 sgRNA | NLS hSaCas9 NLS 3xHA; U6 promoter CXCR4-2 sgRNA | This work |
| 19 | AcrIIA4 | SV40NLS-GGS-AcrIIA4 | Reference [4] |
| 20 | AcrIIC1 | AcrIIC1 | Reference [5] |
| 21 | AcrIIC1 chimera-1 | AcrIIC1 with LOV insertion between Q69 and Y72 | This work |
| 22 | AcrIIC1 chimera-2 | AcrIIC1 with LOV insertion between E68 and Y72 | This work |
| 23 | AcrIIC1 chimera-3 | AcrIIC1 with G-LOV-G insertion behind Y70 | This work |
| 24 | AcrIIC1 chimera-4 | AcrIIC1 with SG-LOV-GS insertion behind Y70 | This work |
| 25 | AcrIIC1 chimera-5 | AcrIIC1 with GSG-LOV-GSG insertion behind Y70 | This work |
| 26 | AcrIIC1 chimera-6 | AcrIIC1 with mCherry insertion between Q69 and Y72 | This work |
| 27 | AcrIIC1 chimera-7 | AcrIIC1 with mCherry insertion between E68 and Y72 | This work |
| 28 | AcrIIC1 chimera-8 | AcrIIC1 with G-mCherry-G insertion behind Y70 | This work |
| 29 | AcrIIC1 chimera-9 | AcrIIC1 with GS-mCherry-GS insertion behind Y70 | This work |
| 30 | AcrIIC1 chimera-10 | AcrIIC1 with GSG-mCherry-GSG insertion behind Y70 | This work |
| 31 | AcrIIC1 chimera-11 | AcrIIC1 with GSG-PDZ-GSG insertion behind Y70 | This work |
| 32 | AcrIIC3 | AcrIIC3 | Reference [5] |
| 33 | AcrIIC1_A48I | AcrIIC1_A48I | This work |
| 34 | AcrIIC1_D15Q | AcrIIC1_D15Q | This work |
| 35 | AcrIIC1_D43F | AcrIIC1_D43F | This work |
| 36 | AcrIIC1_D45F | AcrIIC1_D45F | This work |
| 37 | AcrIIC1_D46E | AcrIIC1_D46E | This work |
| 38 | AcrIIC1_K47Q | AcrIIC1_K47Q | This work |
| 39 | AcrIIC1_M77S | AcrIIC1_M77S | This work |
| 40 | AcrIIC1_N3F | AcrIIC1_N3F | This work |
| 41 | AcrIIC1_N3Y | AcrIIC1_N3Y | This work |
| 42 | AcrIIC1_R36D | AcrIIC1_R36D | This work |
| 43 | AcrIIC1_N3F/A48I | AcrIIC1_N3F/A48I | This work |
| 44 | AcrIIC1_N3F/K47Q | AcrIIC1_N3F/K47Q | This work |
| 45 | AcrIIC1_D15Q/A48I | AcrIIC1_D15Q/A48I | This work |
| 46 | AcrIIC1_N3Y/A48I | AcrIIC1_N3Y/A48I | This work |
| 47 | AcrIIC1_D15Q/K47Q | AcrIIC1_D15Q/K47Q | This work |
| 48 | AcrIIC1_N3F/D15Q | AcrIIC1_N3F/D15Q | This work |
| 49 | AcrIIC1_N3Y/D15Q | AcrIIC1_N3Y/D15Q | This work |
| 50 | AcrIIC1_N3Y/K47Q | AcrIIC1_N3Y/K47Q | This work |
| 51 | AcrIIC1_K47Q/A48I | AcrIIC1_K47Q/A48I | This work |
| 52 | AcrIIC1_N3Y/D15Q/A48I | AcrIIC1_N3Y/D15Q/A48I | This work |
| 53 | AcrX (AcrIIC1_N3F/D15Q/A48I) | AcrIIC1_N3F/D15Q/A48I | This work |
| 54 | AcrX* | AcrIIC1_N3F/D15Q/A48I with GSG-mCherry-GSG insertion behind Y70 | This work |
| 55 | AAV_AcrX | AAV-compatible vector encoding AcrX | This work |

**Supplementary Table 1 | Constructs employed in this study.** NLS, nuclear localization signal.

| Locus | Target sequence (5' to 3') |
|---|---|
| Firefly luciferase | GCGGGGAGAAGGCCAGGGGTCACTCCAG**GATT** |
| IL2RG | CTCTTTCTCCTCAAGGAACAATCAGTG**GATT** |
| NTS4B | GGACAGGAGTCGCCAGAGGCCGGTGGTG**GATT** |
| NTS59 | ACCCCACAGTGGGGCCACTAGGGACAG**GATT** |
| DHFR | GTGATTTTATAGGTAAACAGAATCTGGT**GATT** |
| F8 | GGTTTCTAGTTGTGACAAGAACACTGGT**GATT** |
| EMX1 | GGCCTCCCCAAAGCCTGGCCAGG**GAGT** |
| GRIN2B | GAGAGTAGGCTGGTAGATGGAGTT**GGGT** |
| CCR5 | GGTGGTGACAAGTGTGATCACTT**GGGT** |
| HBB | AGGGTTGCCCATAACAGCATCAG**GAGT** |
| CXCR4-1 | GGACAGGATGACAATACCAGGCA**GGAT** |
| CXCR4-2 | GATGATAATGCAATAGCAGGACA**GGAT** |

**Supplementary Table 2 | Genomic target sites.** The sgRNA-complementary part is underlined. The PAM is shown in bold. Note, the HBB sgRNA contains a 5' G which is not part of the genomic target site.

| Locus | Direction | Sequence (5' to 3') |
|---|---|---|
| IL2RG | fw | ATGACACTGGTGGGTGTTCAG |
| | rv | TCTTCACCTTGCAGGCTCTCT |
| NTS4B | fw | AGAGGAGCCTTCTGACTGCTGCAGA |
| | rv | AGGTCCTGGCCTTGCCTTCGA |
| NTS59 | fw | TGCTTTCTTTGCCTGGACAC |
| | rv | CCTCTCTGGCTCCATCGTAA |
| DHFR | fw | GCAGACTCCACACAGACGGT |
| | rv | GGGCCTACTGAATGATGGTTCAAG |
| F8 | fw | GGGAGAGAACCTCTAACAGAACG |
| | rv | GCTCCAGGTGATGGATCATCAG |
| EMX1 | fw | GGAGCAGCTGGTCAGAGGGG |
| | rv | GGGAAGGGGGACACTGGGGA |
| GRIN2B | fw | AGAATTTTGTAATTGGTTCTACCAAAG |
| | rv | ACAACAGTGGAAGAAAGCTAGGGC |
| CCR5 | fw | GGCAACATGCTGGTCATC |
| | rv | GGTGTAAACTGAGCTTGCTCG |
| HBB | fw | ATGGTGCATCTGACTCCTG |
| | rv | ACTGTACCCTGTTACTTATCCCC |
| CXCR4-1 &-2 | fw | AGAATTTTGTAATTGGTTCTACCAAAG |
| | rv | ACAACAGTGGAAGAAAGCTAGGGC |

**Supplementary Table 3 | Primers for genomic PCRs.** Fw, forward primer; rv, reverse primer.

**Supplementary Note**

To test whether the improved *Nme*Cas9 inhibition observed for the chimeric Acrs was due to blockage of *Nme*Cas9 DNA binding, we performed a luciferase experiment. While d*Nme*Cas9-mediated CRISPR interference (CRISPRi) would provide a simple platform to test for Cas9 DNA binding, the catalytic residues mutated within the HNH domain in d*Nme*Cas9 are critical for AcrIIC1 binding[6]. Therefore, we created two Cas9 expression vectors: The first vector encodes an *Nme*Cas9 nickase (n*Nme*Cas9) bearing a mutated RuvC, but wild-type HNH domain, which mediates reporter knockdown by CRISPRi as well as nicking of the reporter construct. The second vector encodes a d*Nme*Cas9, which exclusively mediates CRISPRi, but cannot be impaired by AcrIIC1. As expected, AcrIIC1 was unable to block CRISPRi by d*Nme*Cas9 (Supplementary Fig. 4). Importantly, while expressing AcrIIC1 rescued reporter expression in the presence of n*Nme*Cas9 (bearing a wild-type HNH and mutated RuvC domain), reporter activity did not exceed that observed in the presence of d*Nme*Cas9 (Supplementary Fig. 4). This indicates that while AcrIIC1 impairs nicking of the reporter, it is unable to block DNA binding required to also impair CRISPRi. In contrast and akin to the AcrIIC3 control, AcrIIC1-mCherry chimera 10 fully rescued reporter expression in the presence of the n*Nme*Cas9 (Supplementary Fig. 4), suggesting that CRISPRi cannot occur in these samples due to impaired *Nme*Cas9 DNA binding.
